# Mass spectrometry-based quantification of proteins and post-translational modifications in dried blood: longitudinal sampling of patients with sepsis in Tanzania

**DOI:** 10.1101/2025.04.22.650109

**Authors:** Matthew W. Foster, Timothy J. McMahon, James S. Ngocho, Asia H. Kipengele, Marlene Violette, Youwei Chen, Deng B. Madut, Robert S. Plumb, A. Ian Wong, Lingye Chen, Grace M. Lee, Philoteus A. Sakasaka, Blandina T. Mmbaga, John A. Crump, Micah T. McClain, Christopher W. Woods, Venance P. Maro, Matthew P. Rubach

## Abstract

The proteomic analysis of blood is routine for disease phenotyping and biomarker development. Whole blood is commonly separated into soluble and cellular fractions. However, this can introduce pre-analytical variability; and analysis of a single component (which is common) may ignore important pathophysiology. We have recently developed methods for the facile processing of dried blood for mass spectrometry-based quantification of the proteome, N-glycoproteome and phosphoproteome. Here, we applied this approach to 38 patients in Tanzania who presented to the hospital with sepsis. Blood was collected on Mitra devices at presentation and 1, 3 and 28-42 days post-enrollment. Processing of 96 total samples was performed in plate-based formats and completed within two days. Approximately 2,000 protein groups and 8,000 post-translational modifications were quantified in 3 LC-MS/MS runs at ∼1.5 hours per sample. Analysis of differential abundance revealed blood proteome signatures of acute phase response and neutrophilic inflammation that partially resolved at the 28-42 day timepoint. Numerous analytes correlated with clinical laboratory values for c-reactive protein and white blood cell counts, as well as the Universal Vital Assessment illness severity score. These datasets serve as proof-of-concept for large scale MS-based (sub)phenotyping of disease using dried blood and are available via the ProteomeXchange consortium (PXD060377).

## INTRODUCTION

The “blood proteome” is a term often applied to studies of cell-free plasma or serum,^1-3^ which can be misleading since it ignores the bulk of total circulating protein contained in erythrocytes and other cell-types. In contrast, the proteomic analysis of whole blood facilitates the use of remote, low-volume and minimally invasive sampling techniques, while potentially minimizing pre-analytical variability associated with plasma separation.^4-7^ Volumetric absorptive microsampling (VAMS) has been increasingly adopted for collection and storage of dried blood and subsequent proteomic and metabolomic analyses.^4,5,8-10^ To increase the breadth of analytes quantified in blood obtained by VAMS, we recently developed methods for serial enrichment and analysis of N-glycopeptides and phosphopeptides from 20 µL of blood captured on a single Mitra device.^11^

Sepsis and related critical illnesses are multi-organ diseases, motivating extensive investigation of the (circulating) plasma proteome,^12-14^ with goals of phenotyping disease severity, predicting mortality and understanding recovery. Standard-of-care management in hospitalized sepsis patients includes routine blood collection for measurement of well-validated proteins and metabolites in plasma or serum.^15^ Analyses of the plasma proteome using remnant clinical samples—which grew exponentially to hundreds of studies during the COVID-19 pandemic— have proven practical and may provide a rapid pathway to the identification of drug targets or for the development of new diagnostics.^16,17^ This contrasts proteomic analyses of dried blood in COVID-19, which is limited to a single study.^6^

Here, we hypothesized that the application of VAMS and comprehensive dried blood analysis in a sepsis cohort would present new opportunities for disease phenotyping. Indeed, the rationale for applying our newly-developed “multi-omic” approach (analysis of proteins, N-glycopeptides and phosphopeptides) is supported by the remodeling of the plasma N-glycoproteome and the peripheral blood mononuclear cell phosphoproteome in sepsis and COVID-19 disease.^18-21^ To demonstrate feasibility of sample collection and analysis, we integrated the collection of dried blood on Mitra devices into an existing biobanking protocol enrolling patients with Sepsis 2 definition (system inflammatory response syndrome features, and suspicion of infection)^22^ presenting to one rural and two urban hospitals in Kilimanjaro, Tanzania.

## MATERIALS AND METHODS

### Subject enrollment and sample collection

Sepsis samples were collected under the approval of the Duke University Health System Institutional Review Board (Pro00101917), the Kilimanjaro Christian Medical University Clinical Research Ethics Committee (No. 2426), and the Tanzanian National Institute for Medical Research (NIMR/HQ/R.8a/Vol.IX/3132). Screening and enrollment criteria are described in **Supplemental Methods**.

To calculate the Universal Vital Assessment (UVA) score,^23^ study clinicians obtained vital signs, oxygen saturation by pulse oximetry and Glasgow Coma score. HIV infection status was ascertained using SD Bioline HIV-1/2 rapid antibody kit, and if positive, this was confirmed with a second rapid antibody test, Uni-Gold (Trinity Biosystems). Complete blood count was performed on EDTA-blood using the Cell-DYN Ruby (Abbott) hematology analyzer. Serum was fractionated from venous whole blood collected into plain red-top tubes, and serum C-reactive protein (CRP) was measured on COBAS Integrate 400-Plus (Roche) chemistry analyzer.

Venous EDTA-blood was collected at enrollment (d0) and at a post-enrollment follow-up visit (with a window of 28-42 days; d28-42). If participants were still hospitalized, collections were also obtained at one day post-enrollment and at three days post-enrollment. After blood draw, tubes were stored at 4-8 °C for up to 3 h and absorbed onto 4 × 20 µL Mitra devices (Trajan) per patient-timepoint followed by air drying in Mitra device containers (“clamshells”) for 2 h. Labels were applied to septum-capped 750 µL Matrix tubes (ThermoFisher), septa were removed and tips dislodged using a fabricated aluminum decapper (**Fig. S1**), stored at -80 °C and shipped on dry ice.

EDTA-blood from four blood type-matched healthy controls was obtained under IRB Pro00094580. Equal volumes were pooled and loaded on Mitra devices dried for 2 h in the presence of desiccant, transferred to Matrix tubes and stored at -80 °C.

### Trypsin digestion

Digestions were performed as previously described.^24,25^ Briefly, under Biosafety Level 2 (BSL-2), 290 µL of 5% w/v sodium deoxycholate (SDC) and 10 mM DTT in 50 mM ammonium bicarbonate (AmBic) were added followed by heating at 80 °C for 30 min on a Thermomixer. At this point, samples were considered “non-infectious”. After cooling for ∼10 min, 15 µL of 400 mM iodoactemide in AmBic was added and samples incubated in the dark at room temp for 30 min. Finally, 15 µL of 200 mM DTT and 20 µL of 12.5 mg/mL TPCK-trypsin (Worthington) in AmBic were added, followed by heating at 37 °C for 2 h. Reactions were quenched by the addition of 38 µL of 20/20/80 (v/v/v) TFA/MeCN/H_2_O (note: a repeater pipette was used to avoid direct contact of the pipette tip with the sample). After vortexing, precipitates were transferred to an Isolute Filter+ plate (Biotage) and filtered into Matrix tubes using a positive pressure manifold (Biotage Pressure+). A study pool QC (SPQC) sample was made by mixing equal volumes of all samples.

### N-glycopeptide enrichment

Hydrophilic interaction chromatography (HIILIC) enrichment was performed as previously described.^26^ Briefly, a 10% (w/v) slurry of Polyhydroxyethyl A (12 µm, 300 Å; PolyLC cat# BMHY1203) in 0.1% TFA (e.g. 2.5 g in 25 mL) was tumbled end-over-end for 30 min. P200 pipette tips were plugged with 3 discs of MK360 (Ahlstrom) and arrayed in a 96- well format, and 200 µL of slurry was added to each tip using a multichannel pipette. Tips were centrifuged at 200 *xg* for 2-3 min per step followed by washing with 2 × 200 µL of 80% (v/v) MeCN:H_2_O containing 0.1% TFA (wash buffer). Next, 80 µL of the digest was diluted with a mixture of 320 µL of MeCN and 4 µL TFA. Samples were passed through tips in two rounds of 200 µL, and this was repeated 3 times. The flow-through was reserved for phosphopeptide enrichment. Tips were washed with 3 × 200 µL wash buffer followed by elution with 2 × 200 µL of 0.1% TFA. An SPQC sample was prepared from equal volumes of all eluents, and 40 µL of each sample was diluted with 160 µL 0.1% FA on Evotips followed by loading and washing with 2 × 50 µL of 0.1% FA.

### Phosphopeptide enrichment

Phosphopeptide enrichment was performed as previously described.^27^ Briefly, 300 µL of HILIC flow-through was adjusted with 100 µL of 20% TFA (v/v) in MeCN and 1 M glycolic acid (GA) followed by enrichment using a Kingfisher Flex (ThermoFisher) using 40 µL of MagReSyn® Ti-IMAC HP (Resyn Biosciences). Enrichment used the manufacturer’s protocol except that 0.25 GA was utilized in binding and wash buffers. Peptides were eluted with 200 µL of 1% NH_4_OH followed by acidification with 50 µL of 10% formic acid. An SPQC was prepared as described previously, and 100 µL of each sample was diluted with 150 µL of 0.1% FA on Evotips and washed with 2 × 50 µL of 0.1% FA.

### LC-MS/MS of unenriched proteome

Matrix tubes were sealed with a Storage Mat III (Costar), and 5 µL of each sample was analyzed by direct injection using a Waters ACQUITY LC with solvent divert as previously described,^24^ with a 1 × 150 mm ACQUITY Premier 1.7 µm CSH C18 column (Waters) at 100 µL/min and gradient of 3-28% MeCN/0.1% FA over 40 min. A Waters Auxiliary Solvent Manager was used to deliver 50% MeCN/0.1% FA at 100 µL/min via the divert valve from 0-2 min and 42-47 min. The LC was interfaced to an Orbitrap Astral MS (ThermoFisher) via the Optamax NG source with heated ESI (HESI) source and with a spray shield installed. An Orbitrap precursor scan used 240,000 resolution from 400-900 m/z with 500% normalized automatic gain control (AGC) target and 50 ms ion transfer time (IT) with a cycle time of 0.6 s. Astral DIA scans used 124 × 4 m/z windows from 400-900 m/z with a 500% AGC, 6 ms IT and 28% normalized collision energy (NCE).

### LC-MS/MS of the N-glycoproteome

Samples on Evotips were analyzed on an Evosep One LC using a 60 sample-per-day method with a Pepsep 8 cm × 150 μm column (1.5 μm particle size) with a PepSep Sprayer and stainless steel (30 μm) emitter. The LC was interfaced to an Exploris 480 and Orbitrap Astral MS using a Nanospray Flex Source. Orbitrap DDA-MS/MS used a 60,000 resolution precursor mass range from 650-2000 m/z, 300% AGC and 25 ms IT, and Orbitrap MS/MS used 30,000 resolution, 200% AGC and auto IT, an isolation window of 1.2 m/z and stepped NCE of 20, 30 and 40%. Cycle time was 1 s.

Astral MS/MS used an Orbitrap precursor scan at 240K resolution from 850-1550 m/z with 300% AGC and 50 ms IT and a 0.6 s cycle time. Astral DDA-MS/MS used an intensity threshold of 1E4, 300% AGC and 10 ms IT, isolation window of 1.2 m/z and NCE of 35%. Astral DIA-MS/MS used 139 × 5 m/z windows from 850-1550 m/z with a 500% AGC, 8 ms IT and 35% NCE.

### LC-MS/MS of the phosphoproteome

Samples on Evotips were analyzed with an Evosep LC and Orbitrap Astral MS using a 60SPD method as described above. The Orbitrap precursor scan used 240,000 resolution from 380-980 m/z with 500% AGC and 50 ms IT with a cycle time of 0.6 s. Astral DIA scans used 149 × 4 m/z windows from 380-980 m/z with a 500% AGC, 6 ms IT and 28% NCE.

### MS Data Analysis

Raw files from unenriched and phospho-enriched datasets were converted to .htrms format and processed in Spectronaut 19 (Biognosys). For analysis of the unenriched proteome, Pulsar searches of DIA data used default settings except for variable protein N-terminal acetylation only and semitryptic N-terminal specificity and used a UniProt reviewed Homo sapiens database (downloaded on 04/22/24) and appended with contaminants and variant sequences (20,434 total entries). Quantification was performed at MS2 level using q-value setting, local normalization and MaxLFQ protein roll-up. For downstream analysis, the data was filtered to remove missing data in replicates of the SPQC samples.

Phosphopeptide-enriched samples were similarly analyzed using trypsin/P specificity and variable pSTY, and quantification used local normalization with modification type filter “Phospho (STY)”, PTM workflow with probability cutoff of 0.75 and 0,^27,28^ and PTM consolidation using linear method. The non-localized PTMsite data (cutoff 0) was filtered in R to remove localized PTM sites that met a probability cutoff of 0.75 in at least 10 samples. Data were further filtered to exclude PTM sites lacking a phospho modification and removal of PTM sites with missing values in SPQC samples.

Glyco-enriched samples acquired by stepped-collision energy (SCE)-DDA acquisition were analyzed using Glyco-Decipher 1.04.^29^ Default settings were used for identification except for semi-trypsin specificity. Quantification used the “match between runs” option. The resulting glycopeptide matrix was filtered to remove features with mass shifts,^30^ followed by quantitle normalization in R. Peptides lacking a N-X(not P)-S/T motif (after extending peptide sequence) were removed and sequences were replaced with motif position. If there were multiple N-X-S/T motifs, both sites were included. Finally, an “Accession” was created with Uniprot protein name, N site and abbreviated glycan composition. Redundant accessions (e.g. reflecting missed cleavages) were tagged with a “1” to “n”, starting with smallest peptide sequence as “1”. The Glyco-N-DIA workflow of FragPipe was used for analysis of Astral DIA data,^31-33^ followed by visualization in Skyline.^34^

### Statistical analysis

Analyses were performed in R-4.4.1 (coding aided by ChatGPT-4o) or ChatGPT-imbedded Python, Graphpad Prism and ggVolcanoR server.^35^ Final figures were made in Inkscape. Multi-omics factor analysis used the MOFA2 R-package^36^ and model training with 6 factors. Additional analyses used k-nearest neighbor imputation (k=5) and log2 transformation prior to statistical analyses. Comparisons between d0 and d28-42 timepoints used paired t-tests with Benjamini-Hochberg FDR correction. Fold change calculations were the average of paired ratios for each individual. Linear regressions used the lm() function in base R followed by calculation of adjusted p-value (“q-value”) by FDR method.

## RESULTS

### Whole blood biobanking and proteomic analysis

Between and May and October 2023, Mitra devices were loaded with venous EDTA-blood collected from subjects enrolled in an ongoing longitudinal sepsis characterization study (**Table S1; Fig. S1**). The total time between collection of the last device and sample processing was approximately 10 months. Ninety-two samples from 38 subjects were randomly distributed across a 96-well plate and interspersed with replicates of a healthy control (HC) pool (**Table S2**). Deoxycholate-assisted in solution digestion was performed on the first day of sample prep, and the next day, 80 µL (∼1 mg) digests were serially enriched for N-glycopeptides using HILIC and for phosphopeptides using Ti-IMAC. A separate study pool QC (SPQC) was made from each sample type (unenriched proteome, N-glycoproteome, phosphoproteome) by mixing equal volumes of all samples.

For analysis of the unenriched blood proteome, we coupled microflow LC (1 mm internal diameter, 100 µL/min)^24,25,37^ to narrow-window Astral data-independent acquisition (DIA),^38,39^ spanning 400-900 m/z to ensure adequate points-per-peak. A 40 min gradient was used to maintain a throughput of 30 samples-per-day, and we used a divert valve to waste the first 2 min and 5 last min of the LC method to minimize source contamination.^24^ Five microliters of each digest (∼60 µg assuming 250 µg/mL protein concentration in blood) were analyzed in singlicate, with four replicates of the SPQC interspersed.

For quantitative analysis of the N-glycoproteome, ∼10% of HILIC eluents were loaded directly onto Evotips and analyzed using a 60SPD Evosep One LC and Orbitrap SCE-DDA method with a 650-2000 m/z precursor selection. For validation and further exploration of quantitative methodologies, DDA and DIA analyses were acquired on the Orbitrap Astral using a NCE of 35%. We adapted a narrow-window glyco-optimized DIA method ^40,41^ with 139 × 5 m/z windows spanning 850-1550 m/z to maximize points-per-peak and provide coverage for the majority of precursors identified in DDA analyses. Finally, for the phosphoproteome analyses, 100 µL (40%) of the acidified Ti-IMAC eluents (and post-enrichment SPQC pools) were loaded onto Evotips and analyzed using a 60SPD LC method and Orbitrap Astral narrow window (4 m/z) DIA.

#### Data analysis summary

A spectral library generated using Spectronaut Pulsar and individual samples identified 34,745 precursors and 2,338 protein groups. Of these, 34,641 precursors and 2,249 protein groups were quantified in at least one sample, with 7% missing protein-group values. The final matrix contained 2,078 protein groups after removing rows with missing values in the SPQC samples. Median percent coefficient of variation (%CV) of protein groups in the preparative replicates of HC samples and technical replicates of SPQC samples were 7.8% and 7.3% respectively.

N-glycopeptide identification and quantification used Glyco-Decipher with “match-between-runs”,^29^ followed by exclusion of features with unannotated mass shifts (e.g. due to “monosaccharide stepping”).^30^ The total matrix had 12,000 quantified glycopeptides across 100 LC-MS/MS runs with 16% missing data; however, after removal of features with missing data in SPQC samples and peptides without an N-X(not P)-S/T motif, there were 4,120 glycopeptides, 7% missing data and median %CV of 31% and 27% in HC and SPQC pools, respectively. Analysis of DIA data—with spectral libraries built from Orbitrap or Astral DDA data—used the glyco-N-DIA workflow in FragPipe.

The phosphopeptide-enriched data was processed in Spectronaut using the PTM workflow and exported at the PTM site level.^27^ For data completeness, we disabled site localization (i.e. probability cutoff 0) for export of quantitative data and filtered the matrix for PTM sites that were localized (probability cutoff >0.75) in ≥10% of samples, followed by removal of non-phosphorylated features and rows that had missing data in SPQC samples. There were 4,430 unique PTM sites with 6.1% missing data and %CVs of 13.7% and 12.3% for HC and SPQC pools, respectively.

### Multi-omics factor analysis identifies acute phase reactants as significant features in all three proteomes

Multi-omics factor analysis (MOFA)^36,42^ was used for integrated analysis of the three datasets, which comprised quantitative data on ∼10,000 features (protein groups, glycopeptides, phosphosites) per sample. The unenriched proteome had the highest weighting of the combination of factors 1 and 2 (**Fig. 1A**), followed by the phosphoproteome. The three proteomes equally weighted factors 2 and 3. The data was first visualized using a “latent factors plot” of factors 1 vs. 2, and 2 vs. 3 (**Fig. 1B-C**). These plots showed expected tight clustering of the SPQC and HC samples, with the most separation between these pools on factor 1 (**Fig. 1B**). Similarly, HC pool and d28-42 samples appeared to cluster more tightly compared to the d0-3 samples along factors 1 and 2.

**Figure 1.**
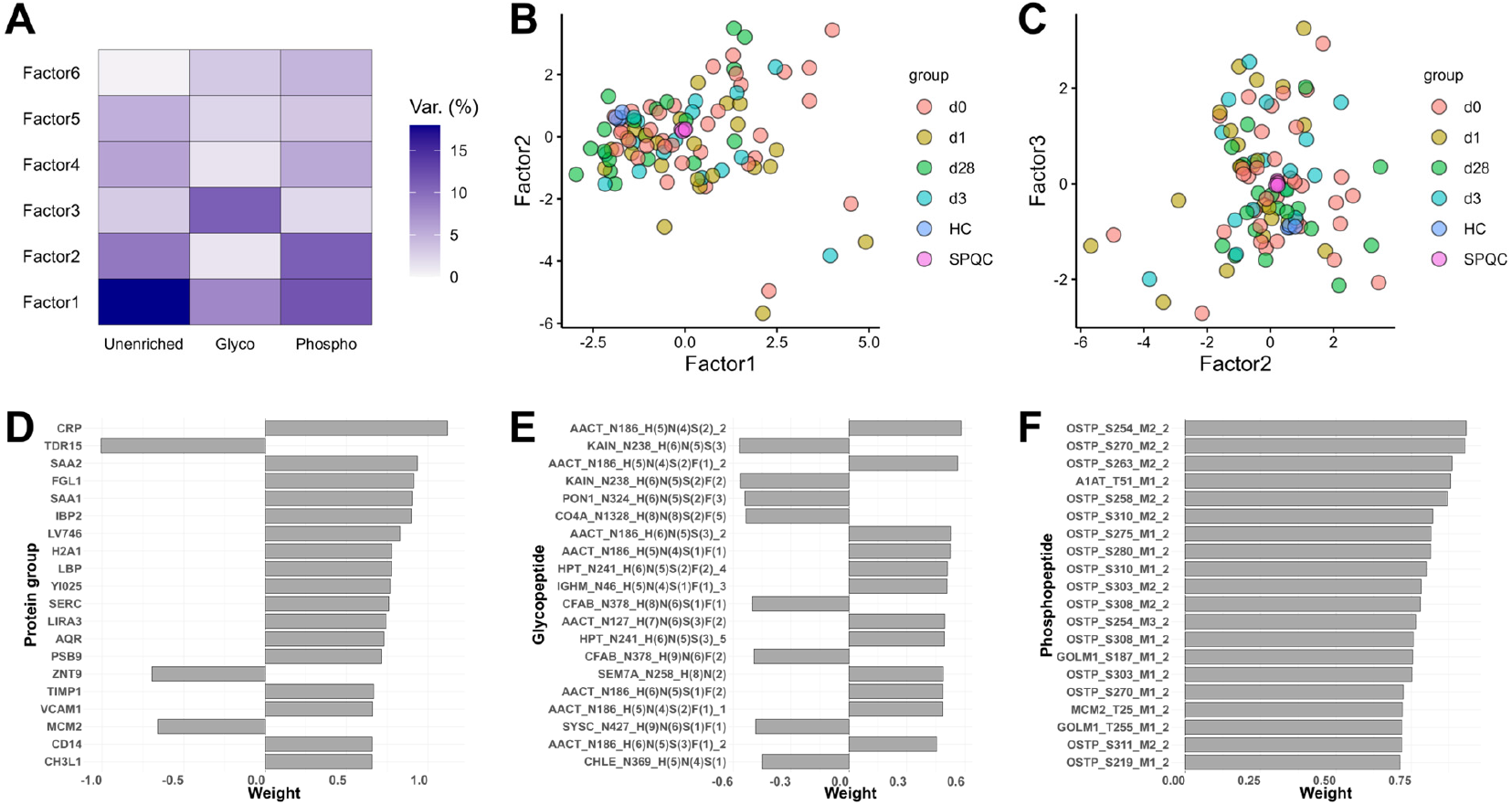
Multi-omics factor analysis of sepsis proteomes. Unenriched, N-glyco- and phospho- proteomes (n=100 each) were analyzed using the MOFA2 R-package. **A**) Variance explained by each proteome within factors 1-6. **B-C**) Latent factor plots of factors 1 vs. 2, and factors 2 vs. 3, respectively. **D-F**) Top 20 features for each proteome weighting factor 1 of the proteome, glycoproteome and phosphoproteome, respectively. Row labels in **D-F**, respectively are: protein name; protein name, glycosylation site and abbreviated glycan composition (H = Hexose, N = HexNAc, S = NeuAc and F = Fucose) and replicate number (for sites represented by more than one peptide due to missed cleavage or semi-specific cleavage); and protein name, PTM site, multiplicity (M1 = one modification, M2 = two modifications) and max number of modifications.

We examined the 20 most-highly weighted features for factor 1 of each proteome (**Fig. 1D-F**). In the unenriched proteome, these included numerous “positive” acute phase proteins (i.e. increasing in infection), including c-reactive protein (CRP), serum amyloids A1 and A2 (SAA1/A2),^43^ lipopolysaccharide-binding protein (LBP) and chitinase 3-like protein 1 (CH3L1; YKL40)^44^ (**Fig. 1D**). Similarly, over half of the most highly-weighted glycopeptides belonged to acute phase reactants, including alpha-anti-chymotrypsin (AACT), paraoxonase 1 (PON1), kallistatin (KAIN) and haptoglobin (HPT), with peptides from the positive acute phase proteins (AACT and HPT) and “negative” acute phase proteins (i.e. decreasing in infection; KAIN and PON1)^45^ having opposite weightings (**Fig. 1E**).

Finally, the secreted phosphoprotein osteopontin (OSTP), which is expressed early in SIRS and sepsis and is diagnostic of severity and prognostic of survival in critically-ill patients,^46-48^ heavily weighted phospho factor 1 (**Fig. 1F**). Two phosphosites from golgi membrane protein 1 (GOLM1)—also known as golgi phosphoprotein 2 or GP73—were are also among the top 20. GOLM1/GP73 is upregulated in viral infections and in the plasma proteome of patients with moderate-to-severe COVID-19,^49,50^ and mouse studies suggest that GP73 is a hormone that links viral infection to blood glucose levels.^50^ Overall, MOFA provided a relatively straightforward approach for data reduction and visualization of the blood proteome, and these results suggested that markers of acute, systemic inflammation contributed to the largest proteome differences between samples in this cohort.

### Acute phase and neutrophil protein signatures dominate recovery in the unenriched proteome

As an initial validation, we compared proteomics data at d0 versus clinical laboratory values for CRP and white blood cells (WBC; **Table S3**). We found a good correlation between the relative values of blood CRP (using un-normalized or normalized maxLFQ values) and serum CRP as measured by COBAS INTEGRA immunoturbidimetric method (**Fig. 2A**). A linear regression analysis using serum CRP as the dependent variable revealed (in addition to CRP measured by MS) significant correlation with group II phospholipase A2 (PA2GA) and other aforementioned acute phase proteins (**Fig. 2B and Table S4**).^51^ A similar correlation was observed between WBC counts and abundant neutrophil proteins or associated mediators (**Fig. 2C**), including the high-mobility group box 1 (HMGB1),^52^ annexin A1 (ANXA1),^53^ histone proteins and integrins. We also performed a similar correlation analysis with the mortality risk UVA score (**Table S4**),^23,54^ and observed a positive coefficient for sepsis markers related to renal dysfunction and tissue injury, including cystatin C (CYTC),^55^ insulin-like growth factor-binding protein 2 (IBP2)^56^ and tenascin (TENA),^57^ as well as a negative coefficient for the negative acute phase proteins kallistatin (KAIN)^58^ and transthyretin (TTHY).

**Figure 2.**
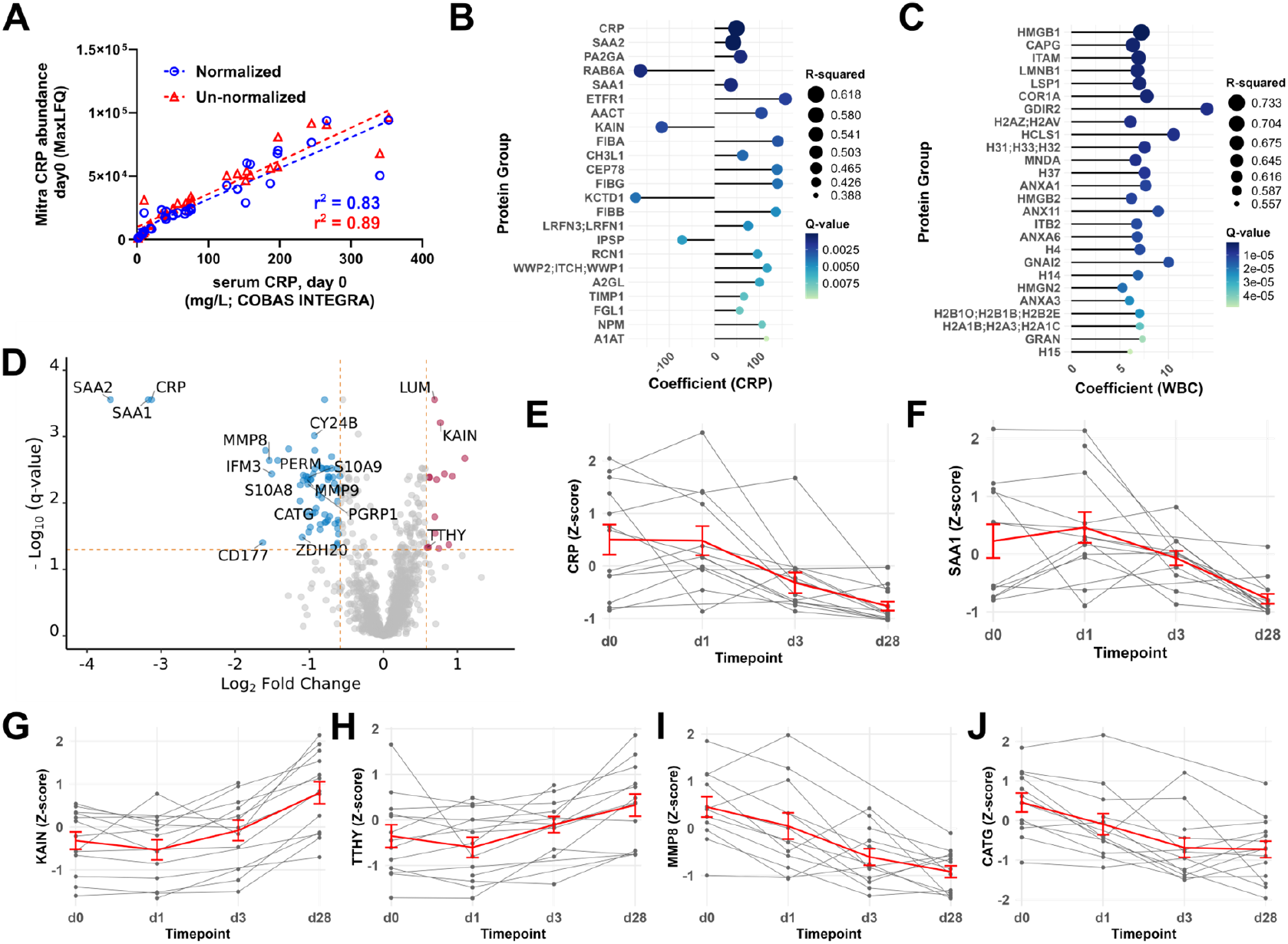
Correlation of blood protein abundance with disease-related clinical values and timepoints. **A**) Correlation of CRP abundance as measured by COBAS INTEGRA immunoturbidimetric method serum (x-axis) versus normalized or un-normalized MaxLFQ abundance from blood proteome (y-axis). Line and r^2^ values were determined by nonlinear least squares regression with GraphPad Prism. **B- C**) Lollipop plot of proteins with the most significant correlation to clinical values of (**B**) CRP and (**C**) WBC at d0 as determined by linear regression. q-value = FDR-adjusted p-value **D**) Volcano plot highlighting the most significant proteome changes between d28-42 vs. 0 (n=19 paired subjects). **E-J**) Trend plots of CRP, SAA1, KAIN, TTHY, MMP8 and CATG protein abundance for n=14 subjects with d0, d28 and at least one other timepoint (n=51 samples total). Data converted to z-score and outliers with z>3 were removed. Individual subject timepoints are in grey, and mean ±s.e.m. is in red.

We further analyzed d0 versus recovery (d28-42) in paired samples (**Fig. 2D and Table S5**). The acute phase proteins (CRP, SAA1/2) had the largest decreases between d28-42 and d0, followed by additional acute phase proteins and numerous abundant neutrophil granule-resident and secreted proteins (S100A8/A9, MMP8/9, CATG, PERM). This signature also included many of the proteins highly-correlated with “hematopoietic” lineage (e.g. ANXA1, PGRP1, ITAM, RETN, CHI3L1 and DEF1) that were identified by Olink proteomics in dried blood spots following COVID- 19 infection.^6^ Statistically significant cell-derived proteins that could be missed in an analysis of plasma alone included: the neutrophil surface receptor CD177, a mediator of integrin signaling that has been suggested as a biomarker of septic shock;^59,60^ cytochrome b-245 heavy chain (CY24B), a subunit of the membrane bound superoxide-generating NADPH oxidase 2 (NOX-2);^61^ IFN-induced transmembrane protein 3 (IFITM3), which has both pro- and anti-viral activity against RNA viruses and regulates platelet reactivity;^62,63^ and the palmitoyltransferase ZDHHC20 (ZCH20), which co-localizes with, palmitoylates and mediates the antiviral activity of IFITM3.^64^ Finally, we examined the longitudinal trends of select proteins in the n=14 subjects that had data for at least three timepoints, including d0 and d28-42. Acute phase proteins like CRP and KAIN tended to be stable between d0 and d1, whereas a steady decrease over time was observed for putative neutrophil-derived proteins such as MMP8 and CATG (**Fig. 2E-I**).

### Phosphoproteome correlations with sepsis severity and inflammatory phenotypes

We next performed similar analyses using the phosphoproteome data. Similar to the MOFA analysis (**Fig. 1E**), many phosphosites of OSTP were positively correlated with UVA score (**Fig. 3A-B and Table S6**). N-terminal histone and Rab44b phosphosites had the strongest correlation with WBC counts at d0 (**Table S6**).

**Figure 3.**
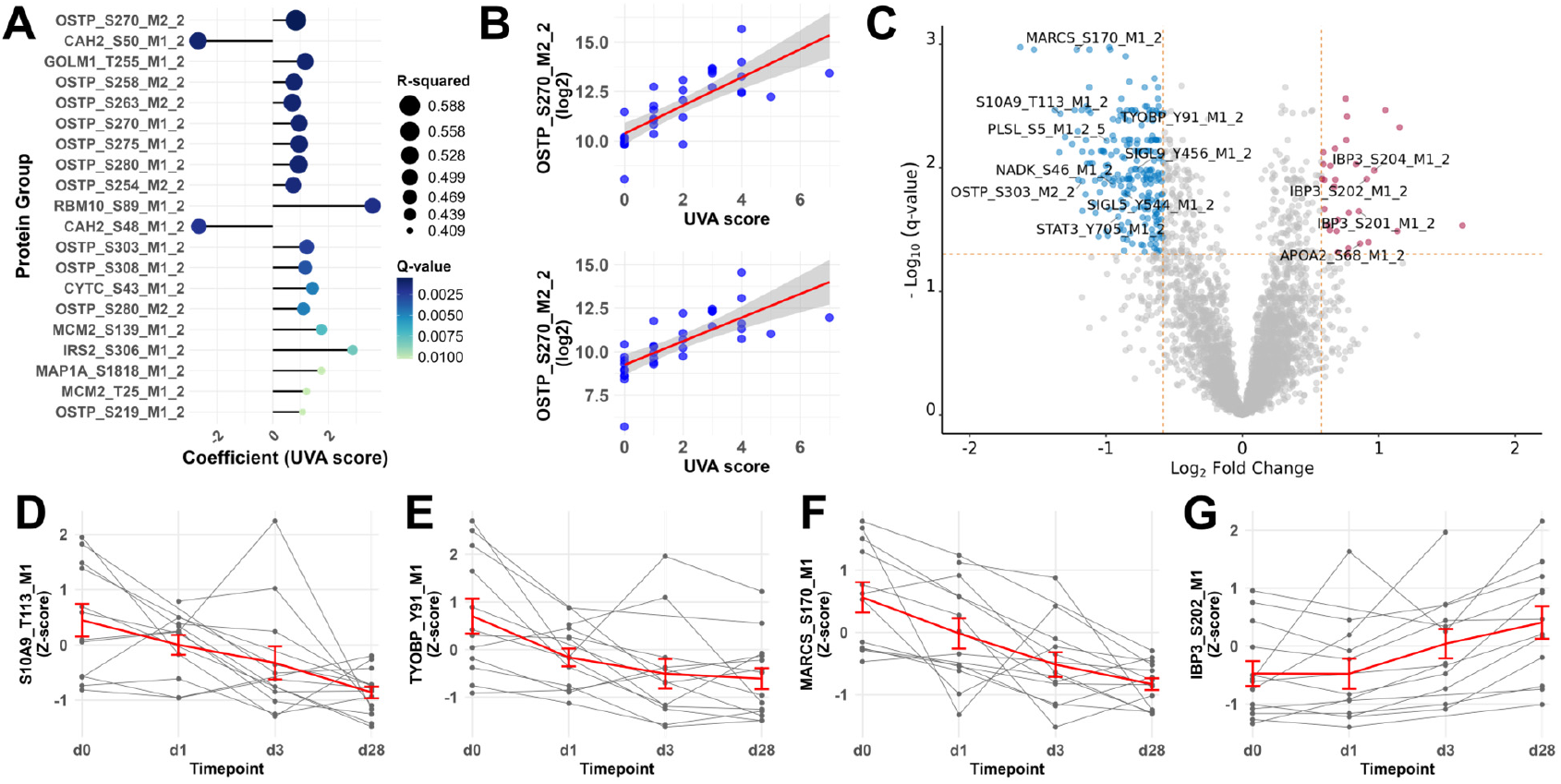
Time- and UVA score-dependent changes in of sepsis severity markers and regulatory phosphosites in the blood phosphoproteome. **A**) Lollipop plot of phosphosites with the most significant correlation to UVA score at d0 as determined by linear regression. **B**) Regression plots for two OSTP phosphosites having significant positive correlation with UVA score, with fitted regression line (red) and 95% confidence interval (gray). The regression line is in red. **C**) Volcano plot highlighting most significant phosphoproteome changes between d28-42 vs. d0 (n=19 paired subjects). **D-G**) Trend plots (as described in **Fig. 1**) for phosphosites (**D**) pThr113 of S10A9, (**E**) pSer170 of MARCS, (**F**) pTyr81 or TYOBP and (**G**) pSer202 of IBP3.

We further compared the d0 vs. d28-42 timepoints using a paired t-test (**Table S7**), which revealed ∼250 phosphosites that had an FDR-corrected p<0.05 and absolute FC>1.5 between presentation to the hospital and recovery (**Fig. 3C**). Phosphosites with roles in neutrophilic inflammation, adhesion and phagocytosis were lower at d28-42, including: Thr113 of S100A9 (S10A9), which regulates the pro-inflammatory activity of secreted S100A8/A9;^65^ Ser5 of l-plastin (PLSL), a target of inflammatory stimulus in leukocytes that modulates integrin-related adhesion;^66^ Ser167 and Ser170 of myristoylated alanine-rich C-kinase substrate (MARCS), which mediates the stimulus-coupled translocation of MARCS from the membrane to cytoplasm;^67^ and stimulatory sites (Ser46/48/64) of NAD kinase (NADK), an enzyme that synthesizes NADPH to support superoxide generation by NOX-2.^68,69^ Regulatory phosphotyrosine sites that were decreased at d28-42 included: the site of activation (Tyr705) of the transcription factor STAT3; Tyr544 of Siglec- 5 (SIGL5/CD170) and Tyr456 of Siglec-9 (SIGL9), which reside in membrane-distal immunoreceptor tyrosine-based inhibitory (ITAM) motifs; as well as the ITAM motif (Tyr91) of the TYRO protein tyrosine kinase-binding protein (TYOBP).^21^ Putative immune cell phosphosites decreased across the timepoints (**Fig. 3D-F**), with insulin-like growth factor binding protein 3 (IBP3) and Apoliprotein A-II (ApoA2), negative acute phase proteins, having the opposite trend (**Fig. 3C&G**).

### N-glycoproteome changes in sepsis and validation with data-independent acquisition analysis

Regression analysis identified a number of N-glycopeptides that correlated with CRP levels and UVA score but much less so with WBC counts (**Table S8**). Interestingly, two distinct clusterin glycopeptides were the most highly correlated with CRP and UVA score, with a negative coefficient, whereas clusterin protein levels were not significantly correlated with either variable (**Fig. S2**). Glycosylated clusterin is reported to be lower in non-survivors in sepsis;^70^ however, on 3 of 96 quantified clusterin glycopeptides correlated with CRP (q<0.05, r^2^>0.4) and 8 correlated with UVA score.

Approximately 10% of the 4,200 quantified glycopeptides were significantly different between d0 and d28-42 (**Fig. 4A**). Of these, 32 corresponding to Asn127 glycoforms of lumican (out of 43 quantified with 0 or 1 missed cleavages; (K)LHINHNN^127^LTESVGPLPK) were higher at d28-42 (**Fig. 4A**). Lumican (LUM) is an extracellular matrix protein that has been shown (in mouse models) to promote the toll-like receptor 4 (TLR4)-dependent innate immune response in bacterial sepsis.^71,72^ We observed a greater effect size for the majority of these glycopeptides peptides than for LUM protein (log2FC of ∼0.7; **Fig. 2D**). Trend plots by subject for LUM and the Hex(5)HexNAc(4)NeuAc(2)Fuc(1) (H5N4S1F1) glycoform both showed the largest increase between d3 and d28-42 (**Fig. 4B-C**). The glycopeptide abundance change was further validated in DIA data using MS2 quantification in Skyline (**Fig. 4D**). We also identified an “off-by-one” error in the GlycoDecipher data (annotation of the M+1 peak of H6N5S1F1 as the M of H5N4S1F3) by manual inspection in Skyline (**Fig. S2**).

**Figure 4.**
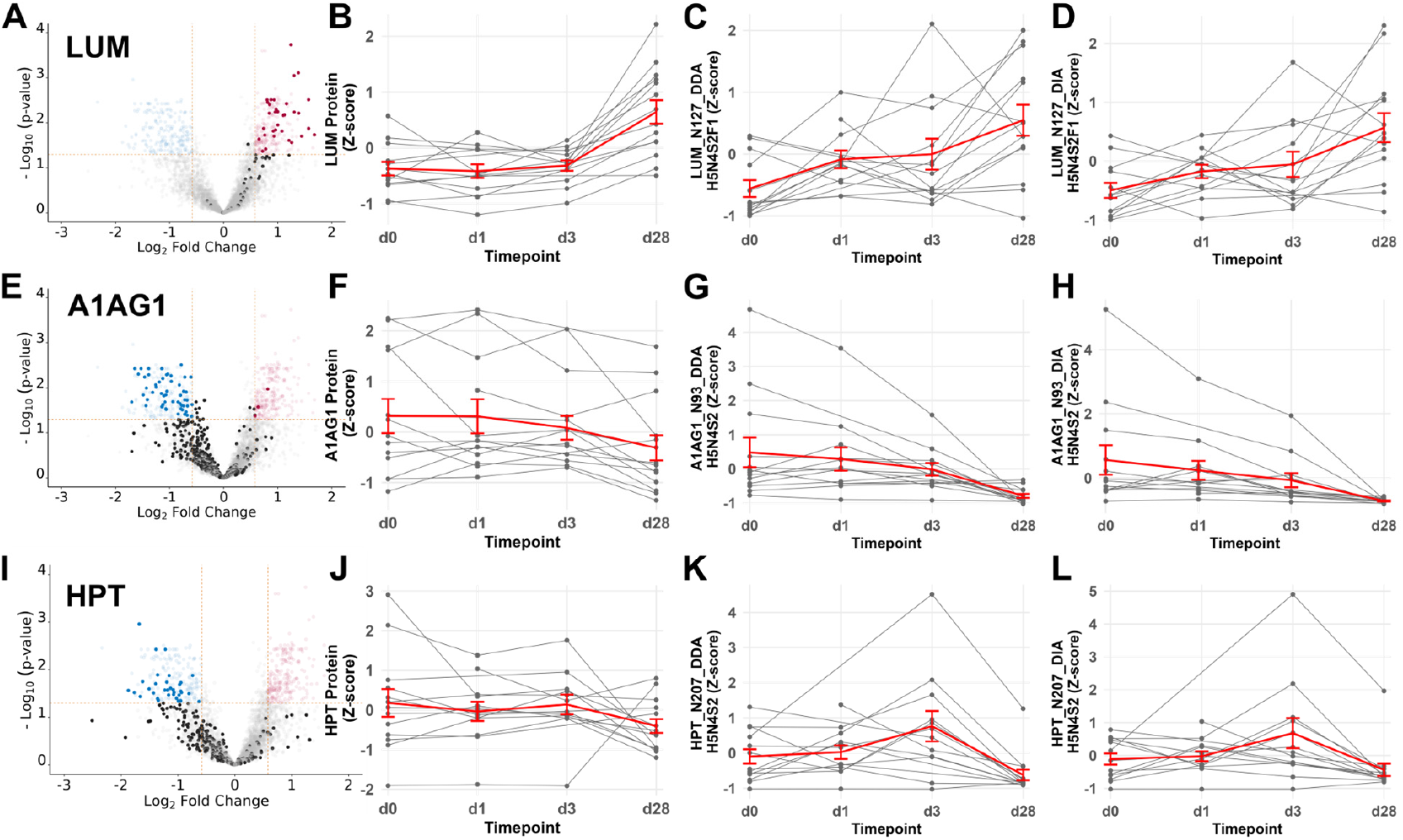
Temporal changes in the blood N-glycoproteome and validation with data-independent acquisition. **A**) Volcano plot highlighting N-glycoproteome changes in lumican (LUM; dark circles) between d28-42 and d0 (n=19 paired subjects). **B-D**) Trend plots (as in **Fig. 1** except that there was no outlier removal) for (**B**) LUM protein, (**C-D**) H5N4S2F1 glycosoform of Asn127 of Lum quantified by MS1/DDA (GlycoDecipher) and MS2/DIA (FragPipe glyco-N-DIA workflow), respectively. **E-G**) and **I-L**) Identical analyses of alpha-1 acid glycoprotein 1 (A1AG1) and haptoglobin (HPT), respectively, as in **A-D**.

There were >400 glycopeptides corresponding to the acute phase protein α-1-acid glycoprotein (A1AG1/Orm1) in the dataset, with many significantly higher at d0 versus d28-42 (**Fig. 4E**). A1AG1 protein was significantly different between d0 and d28-42 but just missed a fold-change cut-off (log2FC ∼-0.57; **Table S9**). Glycoforms at Asn93 have been shown to distinguish bacterial versus viral sepsis.^73^ The H5N4S2 glycoform at N93 of A1AG1 was markedly lower at d28-42 in the DDA/MS1 analysis, with several subjects exhibiting larger changes (**Fig. 4G**); this was replicated in the DIA/MS2 analysis (**Fig. 4H**). Glycosylated haptoglobin (HPT) has received recent attention in sepsis for its role in inflammatory signaling,^74^ and it is represented in the dataset by >250 quantified glycopeptides, many of which were higher at d0 versus d28-42 (**Fig. 4I**). There was a modest but not significant decrease in HPT protein abundance between d0 and d28-42 (**Supplemental Data** and **Fig 4J**). The exemplary H5N4S2 glycoform at Asn207 trended higher between d0 to d3, before decreasing at d28-42, a phenomenon that was replicated between DDA/MS1 and DIA/MS2 datasets (**Fig. 4K-L**).

## DISCUSSION

It is increasingly recognized that dried blood analysis has promise for personalized medicine and disease phenotyping. We have recently focused on the development of methodologies for facile PTM enrichment and proteomic analyses from dried blood collected on Mitra devices.^1175^ Here, we demonstrated that microsampling of whole blood can be easily integrated into established sepsis biobanking protocols and that sample processing and analysis are amenable to repository-scale studies. These methodologies open up opportunities for remote sampling to study longer-term sepsis-related morbidities. While our approach lacks the depth that can be achieved in plasma by enrichment and “next-generation” proteomic approaches,^75^ or by analysis of purified neutrophils or PBMCs,^21,76^ it nonetheless recapitulates many well-described features of sepsis resolution—including reduction in positive acute phase and neutrophil granule proteins—and provides insight into the pathobiology of sepsis that is unlikely to be achieved by analysis of blood plasma or blood-derived cells alone.

Among the three proteomes, we continue to view the N-glycoproteome as the most challenging to analyze and interpret. The future proofing of a high-throughput LC-MS/MS method for increasingly larger scale cohorts is a priority. Of the acquisition types we were able to evaluate here, stepped-collision energy Orbitrap DDA data is, at the moment, the most compatible with existing tools. However, we are encouraged by the analysis of narrow window DIA data using the FragPipe glyco-N-DIA workflow.^40^ By using a consistent sample prep and chromatography approach in future studies, it should be possible to build a comprehensive spectral library for future analyses of this sample type. Skyline (and its integration into the FragPipe workflow) enables real-time QC monitoring of DIA data, facile comparisons of high-resolution MS1- and MS2-level data on identified glycoforms and manual assessment and correction of integration boundaries. Still, we look forward for continued refinement of glycoproteomic workflows and believe that our datasets will be useful for the benchmarking of new and improved analysis tools.

Given the large number of quantifiable red blood cell proteins and PTMs, our analyses to this point belie the potential to discover sepsis-related changes to the erythrocyte proteome. Sepsis is characterized by a reduction in red blood cell deformability,^77^ with phosphorylation of the red blood cell membrane playing an important role in this process.^78^ We measured multiple sites of phosphorylation in the major RBC membrane proteins, including band 3 (B3AT), glycophorins (GLPA/GLPC), ankyrin (ANK1), spectrin (SPTA1/SPTB1) and band 4.1 (4EBP1). Numerous sites in these proteins are sensitive to phosphatase inhibition^79,80^ and are thus expected to be dynamically regulated. However, we did not observe any significant changes to RBC membrane phosphorylation at the acute versus recovery timepoints. It should be noted that we did not quantify the N-terminal pTyr (Tyr8 and Tyr21) regulatory sites in B3AT, and our non-targeted analysis may lack data on other critical sites. It is also possible that the deformability or other phenotypes are poorly reflected in a bulk analysis or more sensitively reflected in molecular changes (e.g. membrane O-glycosylation and lipid content; or RBC metabolites) that will need to be addressed by orthogonal assays.

This study does have several limitations. It lacked a set of matched healthy controls, focused on a relatively mild sepsis cohort and did not fully explore the correlation of ‘omic features with clinical data across the patient-timepoints. We also recognize the need to validate phosphorylation site localization and glycoforms in the dataset. While this can be in part confirmed by manual assignment of MS/MS data, the verification of phosphosite localization may be best accomplished by comparison to authentic standards. Finally, it remains to be determined the extent to which protein abundance changes reflect altered cellularity (e.g. increased blood neutrophils) versus remodeling of granule proteomes,^76^ or similarly whether PTM abundance changes simply correlate with changes in their parent protein.^18,21^ Nonetheless, we believe that this study validates our approach to phenotyping sepsis from dried blood and we hope that it will stimulate future explorations of the “blood” proteome in disease.

## DATA AVILABILITY STATEMENT

Raw and processed data, and metadata, are available at the ProteomeXchange Consortium (PXD060377) via the MassIVE repository (MSV000097000; ftp://massive-ftp.ucsd.edu/v09/MSV000097000/). Skyline results can be viewed or downloaded via the Panorama Public repository (https://panoramaweb.org/SICK_glycoDIA.url).^81^ Code is available at https://github.com/mwfoster/SICK_blood_proteomics.

## ACKNOWLEDGEMENTS

The authors thank Elijah Foster and the Duke University Physics Instrument Shop (Bernie Jelinek and Phil Lewis) for the design and fabrication of the Matrix tube decapper and the FragPipe development team (Alexy Nesvizhskii, Dan Polasky, Fengchao Yu) for assistance and access to a pre-release build of FragPipe 23. This work was supported by funding from the National Instituted of Health, R33-GM146142 (M.W.F and T.J.M.) and R01-AI155733 (M.P.R).

## CONFLICT OF INTEREST STATEMENT

R.S.P is an employee of Waters Corporation. A.I.W holds equity and management roles in Ataia Medical and AI Wong Consulting, LLC. C.W.W. is the chief medical officer for, and owns equity in, Biomeme, Inc.

## Notes

https://proteomecentral.proteomexchange.org/cgi/GetDataset?ID=PXD060377

https://panoramaweb.org/SICK_glycoDIA.url

https://github.com/mwfoster/SICK_blood_proteomics

### Summary of Updates

Updated methods (glycopeptide enrichment) for accuracy and referenced additional pre-print for method development.

