## Supplemental Methods for "Mass spectrometry-based quantification of proteins and post-translational modifications in dried blood: longitudinal sampling of patients with sepsis in Tanzania"

**Inclusion and eligibility.** Screening and enrollment were conducted Monday-Friday from 8 am to 4 pm in the medical wards, emergency departments and outpatient clinics of three hospitals in the Kilimanjaro Region of Tanzania: St Joseph's Council Designated Hospital, Kibosh Council Designated Hospital, and Mawenzi Regional Referral Hospital. Study staff screened all patients ≥ 10 years of age presenting with an acute complaint. Eligibility was based on a modified SEPSIS-2 definition using the following criteria: at least two of the vital sign abnormalities defined by SIRS (tympanic temperature > 38 °C or < 36 °C, heart rate > 90 beats per min, respiratory rate > 20 breaths per min) plus infection suspected as the cause of illness. Eligible patients were excluded if they were a prisoner or refugee, unable to speak English or Kiswahili, unable to provide consent (and without an appropriate representative for proxy consent) or deemed not to have an infectious cause of illness by the treating clinician. Eligible participants or their authorized representatives provided written informed consent prior to sample collection.

**SUPPLEMENTAL FIGURES**

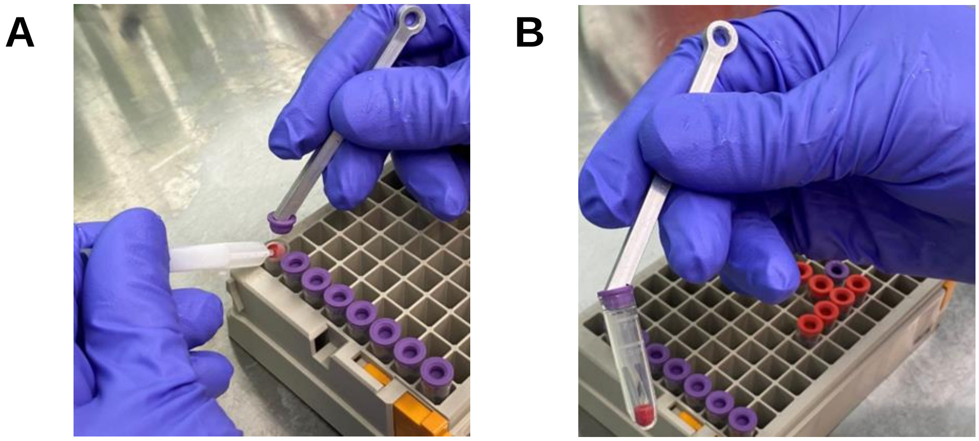


**Figure S1. Storage of Mitra "tips" in Matrix tubes. A)** The septum cap of a 0.75 mL Matrix Tube (ThermoFisher) was removed using a custom fabricated aluminum decapper and the absorptive tip of a Mitra device was plated just inside the edge of the tube. **B)** Re-capping the Matrix tube dislodged the Mitra tip and provided an air-tight storage solution.

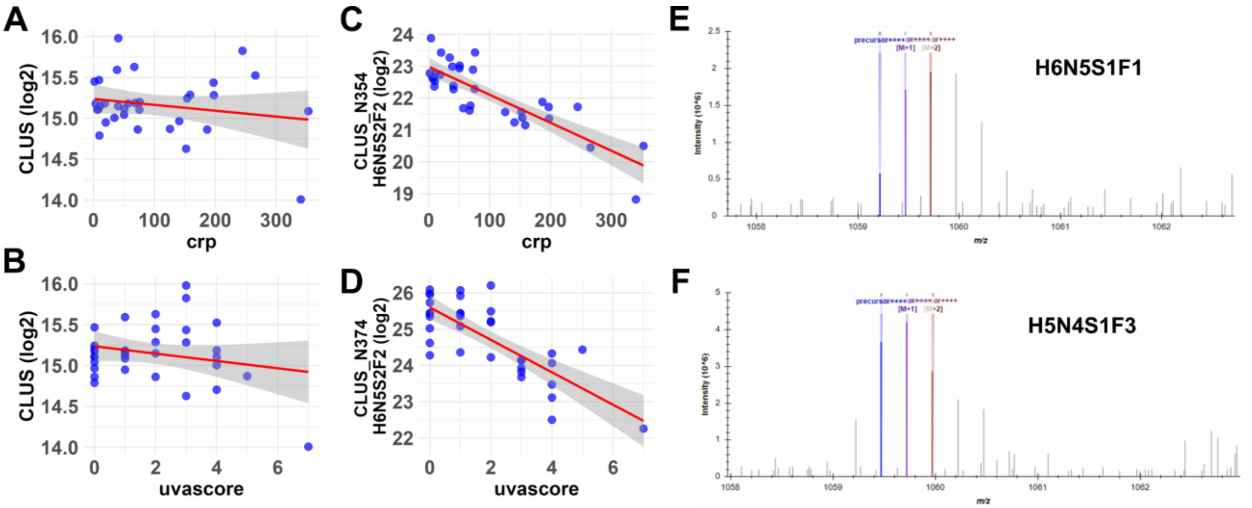


**Figure S2. N-glycopeptide correlation with clinical data, and validation of glycopeptide identifications A-D**) Regression plots for clusterin protein with (**A**) CRP and (**B**) UVA score as dependent variables; and clusterin glycopeptides that had significant correlation with (**C**) CRP and (**D**) UVA score, with fitted regression line (red) and 95% confidence interval (gray). **E-F**) Precursor spectrum from Skyline with M, M+1 and M+2 peaks labeled for Lumican Asn129 glycopeptide with H6N5S1F1 (+2350.8) or H5N4S1F3 (+2351.9) identified by GlycoDecipher. The latter is an example of an "off-by-one" error.
